# Expected impacts of climate change on tree ferns distribution and diversity patterns in subtropical Atlantic Forest

**DOI:** 10.1101/2020.01.16.909614

**Authors:** André Luís de Gasper, Guilherme Salgado Grittz, Carlos Henrique Russi, Carlos Eduardo Schwartz, Arthur Vinicius Rodrigues

## Abstract

Tree ferns are common elements in the Atlantic Forest domain, sometimes reaching more than half of total dominance at forest sites. Just as most groups, climate change might impact the distribution and diversity of tree ferns. To investigate the extent of these impacts in the subtropical Atlantic Rainforest, we measured the changes in species distribution, α- and β-diversity between current climate and future climatic scenarios for 2050. Most tree ferns species tend to lose their distribution area. Hence, species richness tends to decrease in the future, especially in the Rainforest sites. In general, β-diversity tend to not change on the regional scale, but some sites can change its relative singularity in composition. Our results show that climate change can impact distribution and α-diversity of tree ferns, but with no trend to cause homogenization in the tree ferns of the study area. Protected Areas (PAs) in our study region manage to withhold more α-diversity than areas without PAs — the same applies to β-diversity. Our study offers a new light into the effects of climate change in tree ferns by integrating the evaluation of its impacts on distribution, α- and β-diversity in all study areas and inside PAs.

## INTRODUCTION

Tree ferns are expressive elements in (sub)tropical forest formations (Tryon & Tryon 1982), sometimes establishing monodominant forests (Schwartz & Gasper 2020). Consequently, tree ferns take a significant part in the dynamics of ecosystems and may affect the regeneration of woody species and nutrient cycling (Brock *et al.* 2016). Besides, they contribute to ecological succession (Arens & Baracaldo 1998), biomass stock in tropical forests (Sarmiento *et al.* 2005), and provide microhabitat for several epiphytic plants — many of them occurring exclusively on tree ferns’ caudices (Wagner *et al.* 2015).

In tropical forests, tree ferns have suffered intense exploitation due to the ornamental use of their caudices (Hoshizaki & Moran 2001; Eleutério & Pérez-Salicrup 2006), causing populational exhaustion of many species (Santiago *et al.* 2013). Combined with this, the actual fragmentation in the Atlantic Forest (Ribeiro *et al.* 2009) plus climate change (IPCC 2014; Lima *et al.* 2019) are other potential sources of threat to tree ferns. Together, these threats might influence the density and distribution of these species as well as the locations of suitable areas to grow and reproduce.

Moreover, despite tree ferns being an important group in forests’ structure, these plants are historically neglected in floristic and ecological studies (Weigand & Lehnert 2016). The main families of tree ferns in the subtropical Atlantic Forest are Cyatheaceae and Dicksoniaceae. The former (about 15 species) exhibit a preference for warm, humid, and low seasonal climates (Bystriakova *et al.* 2011). On the other hand, the latter is represented by *Dicksonia sellowiana*, a species that inhabits higher and colder environments (Gasper *et al.* 2011), and *Lophosoria quadripinnata*, a species that grows in ravines, doing best on moist, well-drained soils and in full sun until 2,000 m in eastern Brazil (Lehnert & Kessler 2018).

Ferns richness is associated with water availability (Kessler *et al.* 2011), and rainfall regimes modifications could impact ferns distribution. So, the predicted shifts in rainfall and temperature in the subtropical Atlantic Forest, projected by the Intergovernmental Panel on Climate Change (IPCC) should impact these plants. Therefore, a reduction of colder environments and increasing of warmer and moister environments may impact temperate species (VanDerWal *et al.* 2013) as well as *D.sellowiana* — an already endangered species — through the restriction of its occurrence area.

In this regard, our study sought to predict the impacts of future climate changes in α- and β-diversity of tree ferns in the subtropical Atlantic Forest, as well as to predict the impacts in the potential distribution of each species. Our first hypothesis (H_1_) is that species from both families will change their potential distribution areas. We expect Dicksoniaceae species will have their potential distribution area reduced (especially *Dicksonia sellowiana*) because of their association with colder habitats and Cyatheaceae species will increase their potential distribution areas since they generally occur along hot and humid regions. In conjunction — and as a consequence of — with these changes in species distribution, we also expect changes in α- and β-diversity will also change (H_2_). Since we predict that Cyatheaceae species will increase their distribution range, hence increasing the overlap in species areas, we predict an increase in regional α-diversity and a decrease in β-diversity— i.e., less variation in species composition among sites, leading to a homogenization of our study region.

## MATERIAL AND METHODS

### Study area

The study area is delimited by the subtropical Atlantic Forest, floristically distinct from the tropical Atlantic Forest (Eisenlohr & Oliveira-Filho 2014). The subtropical Atlantic Forest occurs in the southernmost portion of Brazil as well as in parts of Argentina and Paraguay (Galindo-Leal & Câmara 2005). The predominant climate type is Cfa (temperate humid with hot summer), with some areas fluctuating to Cfb (temperate humid with temperate summer) (Alvares *et al.* 2013). The relief ranges from sea level to altitudes near 1,200 m, including peaks that reach almost 1,900 m. Distinct forest types can be found, which includes “Restinga”, on the coastal areas; Rainforests, in low altitudes at the coastal region (< 800–900 m); Mixed Forests (*Araucaria* forest), generally in areas with altitudes 800 m a.s.l.; and Semideciduous Forests in low areas with high seasonality in precipitation (Figure 1).

**Figure 1.**
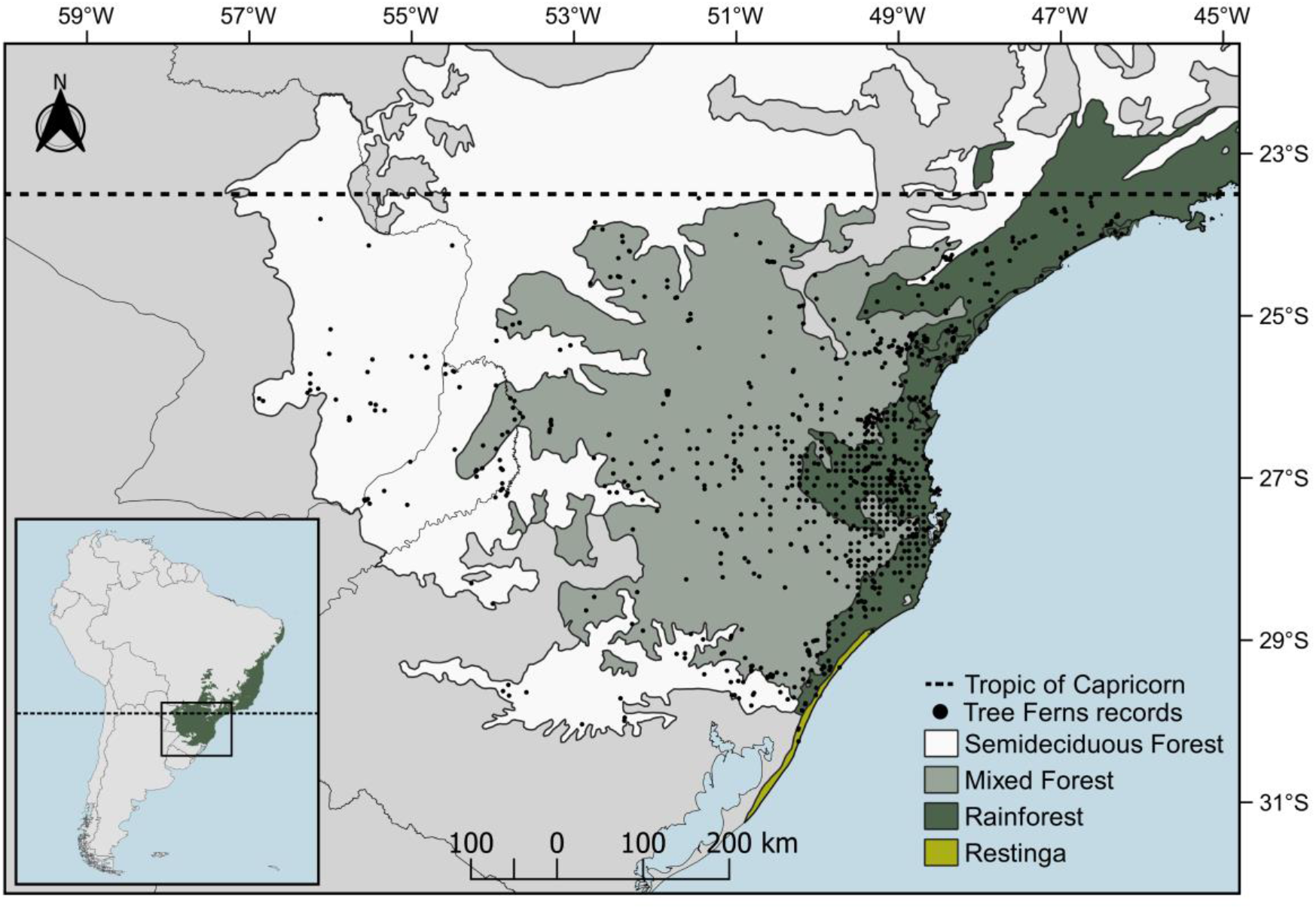
Delimited study area and vegetation types of subtropical Atlantic Forest. Black dots represent locations (plots) of tree ferns’ records used in the study.

### Data gathering

Among the species that occur in subtropical Atlantic Forest (following Brazilian Flora 2020 in construction, 2020; Fernandes, 1997; Weigand & Lehnert, 2016), we identified 1,472 valid records and 15 species from the literature (Fernandes 1997; Vibrans *et al.* 2010) and digital herbaria (using SpeciesLink – http://splink.cria.org and GBIF — http://gbif.org). Distributions outside of those reported in the literature were checked and, if there was any doubt about the record, we removed it from the analysis. Due to its low number of registers, *Cyathea uleana* was removed from the analysis. The occurrence records, as well as their respective references, are available in the GitHub repository (https://github.com/botanica-furb/treeferns).

### Climatic data

Climate data (future and present) were obtained from two high-quality data sets: CHELSA (Karger *et al.* 2017), based on the ERA-Interim climatic reanalysis (Dee *et al.* 2011), and WorldClim v2.1 (Fick & Hijmans 2017), based on spatially interpolated climate data from meteorological stations. Data for 2050 (2040–2060) were based on two IPCC scenarios: “optimistic” (WorldClim SSP1-2.6 and CHELSA RCP2.6) and “pessimistic” (WorldClim SSP5-8.5 and CHELSA RCP8.5). Briefly, the optimistic scenario assumes net negative CO_2_ emissions after 2020 (0 by 2100), and the pessimistic assumes a “business-as-usual” scenario (IPCC, 2014).

For future projections with CHELSA, the ten following General Circulation Models (GCMs) were selected: CanESM2 (Arora *et al.* 2011), CESM1-CAM5 (Hurrell *et al.* 2013), CNRM-CM5 (Voldoire *et al.* 2013), FGOALS-g2 (Li *et al.* 2013), GFDL-ESM2G (Dunne *et al.* 2012), HadGEM2-AO (Martin *et al.* 2011), MIROC-ESM (Watanabe *et al.* 2011), MPI-ESM-MR (Giorgetta *et al.* 2013), MRI-CGCM3 (Yukimoto *et al.* 2012), and NorESM1-M (Bentsen *et al.* 2013). To project future scenarios with WorldClim v2.1 we used six GCMs: BCC-CSM2-MR (Wu *et al.* 2019) CanESM5 (Swart *et al.* 2019), MRI-ESM2 (Yukimoto *et al.* 2012) CNRM-CM6-1 (Voldoire *et al.* 2019), IPSL-CM6A-LR (Boucher *et al.* 2020) and MIROC6 (Tatebe *et al.* 2019). All the GCMs above were chosen for being the most interdependent in both data sets, following Sanderson *et al*. (2015). In this sense, they encompass the highest amount of uncertainty in the future climate. The modeling and projection to future scenarios were made using 30 arc sec resolution for CHELSA and 2.5 arc minutes for WorldClim, — since the latter does not have future data at 30 arc sec resolution yet. To reduce the collinearity between the chosen 19 bioclimatic variables (BIOCLIM 1–19) we used the Variance Inflation Factor (VIF) via *vifstep* function on *usdm* package (Naimi 2015; R Core Team 2020). This function generates 5,000 random points across the climate layers and calculates the VIF for each variable. We used this step-by-step procedure to obtain a set of weakly correlated variables. At each step, the layer with the highest VIF is removed from the set. The process is repeated until only variables with a VIF below 5 remains. This procedure reduced the 19 climate variables to seven in the CHELSA data set — Annual Mean Temperature (BIO01), Isothermality (BIO03), Temperature Seasonality (BIO04), Mean Temperature of Wettest Quarter (BIO08), Mean Temperature of Driest Quarter (BIO09), Annual Precipitation (BIO12), and Precipitation Seasonality (BIO15) — and six in the WorldClim data set — Isothermality (BIO03), Temperature Annual Range (BIO07), Mean Temperature of Wettest Quarter (BIO08), Mean Temperature of Driest Quarter (BIO09), Precipitation of Wettest Month (BIO13), and Precipitation of Coldest Quarter (BIO19).

### Species distribution modeling

The systematic sampling effort of the Floristic and Forest Inventory of Santa Catarina (IFFSC; Vibrans *et al.* 2020) created a sampling bias in the major portion of our records locations. This bias can be identified in Figure 1, where a gridded pattern of equally distant points (10 km at most) occurs over Santa Catarina’s territory. Thus, to spatially filter the registers (Fourcade *et al.* 2014), a buffer of 0.2 degrees (≈20.5 km) was defined as the minimum distance between occurrence records using the *ecospat* package (Di Cola *et al.* 2017). This distance was enough to disaggregate clustered records.

To model each species distribution, we used MaxEnt (Phillips *et al.* 2006) — with cloglog outputs (Phillips *et al.* 2017) and 10,000 background points (Phillips *et al.* 2009). We opt to model species distribution using a single ‘tuned’ algorithm since there are no particular benefits of using an ensemble (Hao *et al.* 2020) — and MaxEnt alone performs just as well as an ensemble (Kaky *et al.* 2020). Despite being a high-performance algorithm, some studies have already shown that MaxEnt’s default settings can perform poorly (Warren & Seifert 2011; Radosavljevic & Anderson 2014; Warren *et al.* 2014). This occurs because MaxEnt automatically chooses its feature classes (FC) based on the sample size and the regularization multiplier (RM), a penalization parameter, is fixed at 1.

Thus, to maximize the predictive performance (Warren & Seifert 2011) it is recommended to (a) generate multiple combinations of RM and FC; and (b) select the best model out of the RM × FC combinations using Akaike’s Information Criteria corrected for small sample sizes (AICc, Burnham & Anderson 2002). In this manner, for each species (13) in each climate data set (separately), we generated 63 models based on the combination of nine RM values (0.5 to 5, increasing by 0.5) and seven combinations of FC (L, H, LQ, LQH, LQP, LQHP, LQHPT). From the set of 63 available models for each species, the one with the lowest value of ΔAICc was selected as the optimal.

The discrimination performance of each model selected by AICc was done using the Area Under the Curve (AUC) of the receiver operating characteristic curve (ROC; Fielding & Bell, 1997), based on a geographically structured cross-validation procedure (Radosavljevic & Anderson 2014). This procedure partitions as equally as possible the presence and background points into five spatial bins. The training is conducted in four bins and the remaining one is used for testing (Muscarella *et al.* 2014). This partitioning method (block) is known for providing the best spatial independence between training and testing data sets (Radosavljevic & Anderson 2014) and providing the best transferability of models across space and time (Fourcade *et al.* 2018). The best model for each species, selected via AICc, was projected into two different future scenarios (optimistic and pessimistic) represented by different GCMs predictions (10 in CHELSA, 6 in WorldClim) in each one of them. We did not ensemble the GCMs to preserve the intrinsic variability among them. Thus, we can account for the uncertainty in the future climate (Porfirio *et al.* 2014), and hence the uncertainty in the change in species distribution and diversity. The modeling procedure above was conducted using *ENMeval* (Muscarella et al. 2014) package.

Finally, we used a site-specific approach to turn the continuous maps generated by MaxEnt into binary maps (presence or absence). The *pS-SDM+PRR* method (Scherrer *et al.* 2018) consists of first summing the continuous probabilities of each species occurring in each site to obtain the predicted species richness. Then, the species are ranked in decreasing order of probability of occurrence in each site. Following the ranking, the best-classified species are selected until the predicted species richness is obtained. Species not chosen in the last step are considered absent in the site.

### Diversity metrics

To understand how climate change will affect diversity patterns, the study area was divided into 2521 hexagonal cells of ≅ 195 km^2^ (±53), not always regular due to the shape of our study region. All binary SDM predictions were rescaled to this hexagon cells for diversity analysis. A species was considered present in a cell if MaxEnt’s prediction covered more than 25% of the hexagon area. Then, based on scaled species distribution in the hexagon cells, we measured for each scenario α- and β-diversity.

We measured α-diversity as the number of species present in each cell, i.e., species richness. β-diversity was calculated using two indices: total β-diversity (BD_TOTAL_) and Local Contribution to β-Diversity (LCBD; Legendre & De Cáceres 2013). BD_TOTAL_ is calculated as the total variance of the site-by-species community matrix and LCBD is the relative contribution of a site to the total variation (Legendre & De Cáceres 2013). Thus, LCBD represents the compositional singularity of each hexagonal cell. Before conducting the calculations above, the community matrix was first Hellinger-transformed. All β-diversity metrics were calculated using *adespatial* (Dray *et al.* 2019) and the functions *beta.div*. All mentioned analyses were conducted in R (R Core Team 2020).

### Evaluating changes in distribution and diversity

We used two basic statistic metrics to evaluate changes in species distributions, α-, and β-diversity in each future scenario: 1) mean differences; and 2) confidence intervals (CI; at 95%) based on the differences between current and future maps. In each future scenario (optimistic, pessimistic) predicted by each GCM – 10 in CHELSA and 6 in WorldClim – we measured the difference between each GCM prediction and current values for each metric of interest. From these values of difference, we calculated the mean and CI. The upper and lower CIs were calculated as 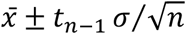, where 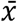 is the sample mean; *t*_*n*−1_ is the critical t-value with *n*– 1 degrees of freedom and area of *α*/2 in each tail; *σ* is the standard deviation; and *n* is the sample size. We decided to use a Student’s t-distribution since it is a better fit for small sample sizes (10 and 6). Hence, we can obtain a significant α at 0.05 for a standard t-distribution.

To evaluate the changes in the species distribution area, we calculated (Fa − Ca/Ca) × 100, where Fa is the area of distribution in the future climate scenario, and Ca is the area of distribution in the current climate scenario. This equation returns the changes in areas as a percentage: positive values indicate an increase in species area, while negative values indicate a decrease. We calculated these changes in area based on the number of cells that each species was predicted to occur in the resolution of each environmental data source, that is, 30 arc sec in CHELSA and 2.5 arc minutes in WorldClim. To assess the changes in α-diversity we measured the differences in richness in each one of the 2521 hexagonal cells and calculated the mean and CI as described above. Following, we built maps that show mean differences for sites in which the range of CI does not include 0 (i.e, the change is statistically significant). The same procedure was used to evaluate the changes in LCBD. On a regional scale, we measured the changes in α-diversity calculating differences of the mean richness of sites between each future GCMs predictions and current maps. Then, the mean and CI were calculated from the differences in mean richness. Changes in total β-diversity were also calculated as mean differences between future GCMs predictions and current maps.

Finally, we evaluated α- and β-diversity changes on a regional scale using only grid cells that overlap Protected Areas (PAs) in the study region — applying the same procedure described above. Boundaries of PAs were obtained from the World Database on Protected Areas (UNEP-WCMC 2019). Therefore, we were able to compare the change in diversity indices between protected and non-protected areas in any specific scenario.

## Results

### Species distribution

The predicted impacts of climate change on species distribution were similar in both climate data sets and future scenarios (Table 1; Figure S01-13). The changes inside PAs were also similar (Figure S13-14). However, some species differ in behavior between CHELSA and WorldClim: *Sphaeropteris gardneri* more than doubled its distribution in CHELSA’s optimistic scenarios while reduced its distribution in WorldClim; *Cyathea atrovirens* and *C. delgadii* suffered a reduction in distribution in CHELSA while an increment in distribution occurred in WorldClim. These differences, nonetheless, only happened outside areas within PAs. In PAs, for example, if a species distribution was expected to decline in either climate data set or scenario the same pattern arises in the other. Taking into consideration all possible combinations (104) of climate data sets (2) × region (2) × scenarios (2) × species (13), only five species of Cyatheaceae had the potential to increase its distribution. Both Dicksoniaceae species are predicted to lose suitable areas. Individual maps of species distribution can be seen in the supplementary material (Figure S01-13) and online in a Shiny interface (https://avrodrigues.shinyapps.io/tferns).

**Table 1.**
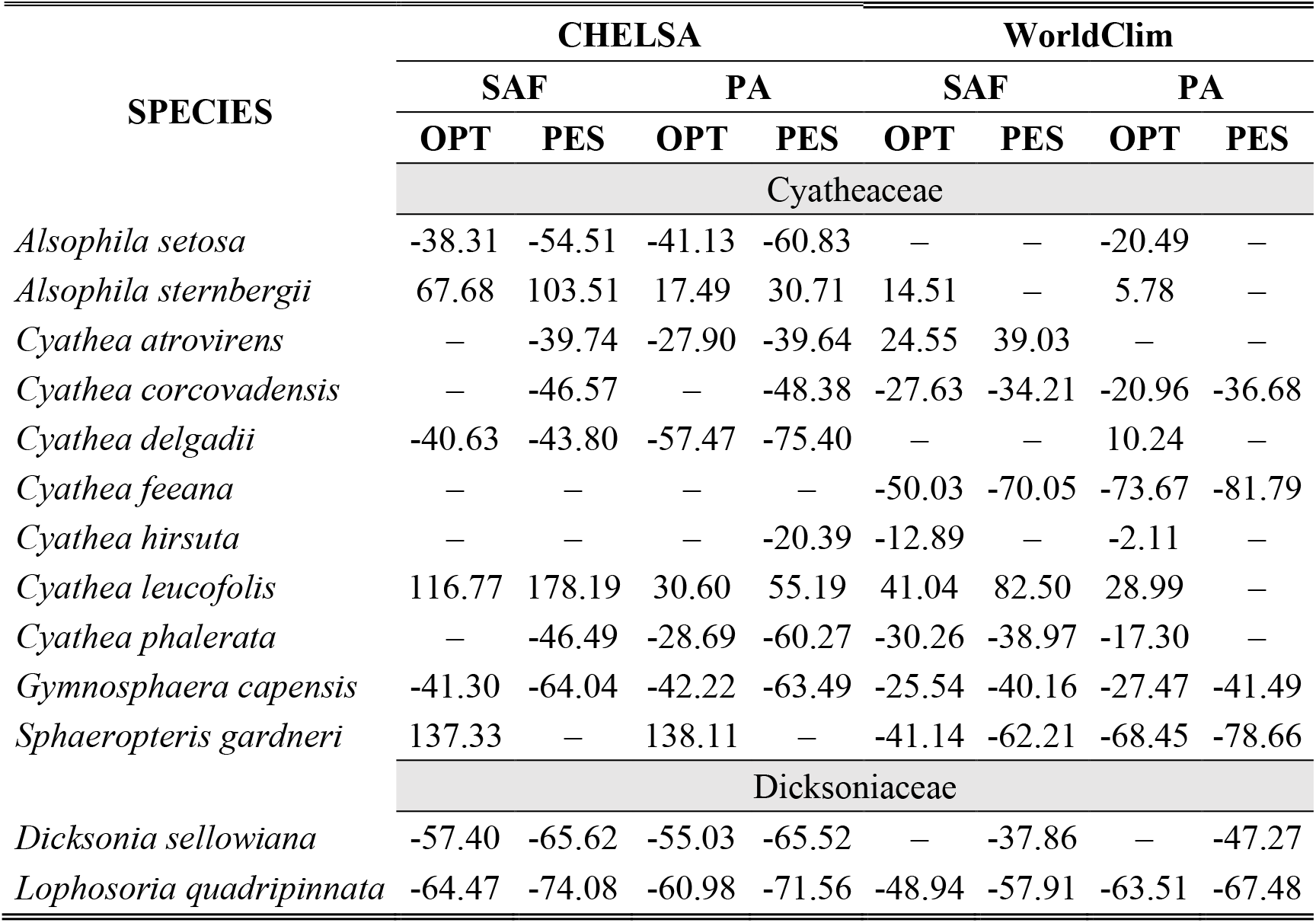
Mean changes (%) in the distribution of species regarding the climatic data set, future scenario, and region. Negative values represent predicted loss in species area and positive values represent predicted gains. Omitted values are non-significant changes. SAF – All subtropical Atlantic Forest; PA – Only Protected Areas; OPT – optimistic future scenario; PES – pessimistic future scenario.

### Diversity patterns

#### α-diversity

In CHELSA, the highest α-diversity values (current and future scenarios, Figure 2; https://avrodrigues.shinyapps.io/tferns) were found in the Atlantic Rainforest areas. The same pattern was found in WorldClim. In addition, WorldClim also exhibited elevated richness in a transitional zone between *Araucaria* forest and Semideciduous forest in Rio Grande do Sul, Brazil. In the Mixed and Semideciduous Forest, where α-diversity is lower, few changes are observed between climate data sets and scenarios. The current regional mean richness in CHELSA is 3.99 (sd: ± 2.66) in all subtropical region and 5.78 (sd: ± 3.27) in areas within PAs. In WorldClim these values are 4.37 (sd: ± 3.12) and 6.64 (sd: ± 3.76), respectively. A reduction in the regional mean richness occurs in both future scenarios and climate data sets, being more intense in the optimistic scenario (Table 2). The same pattern replicates inside PAs.

**Table 2.**
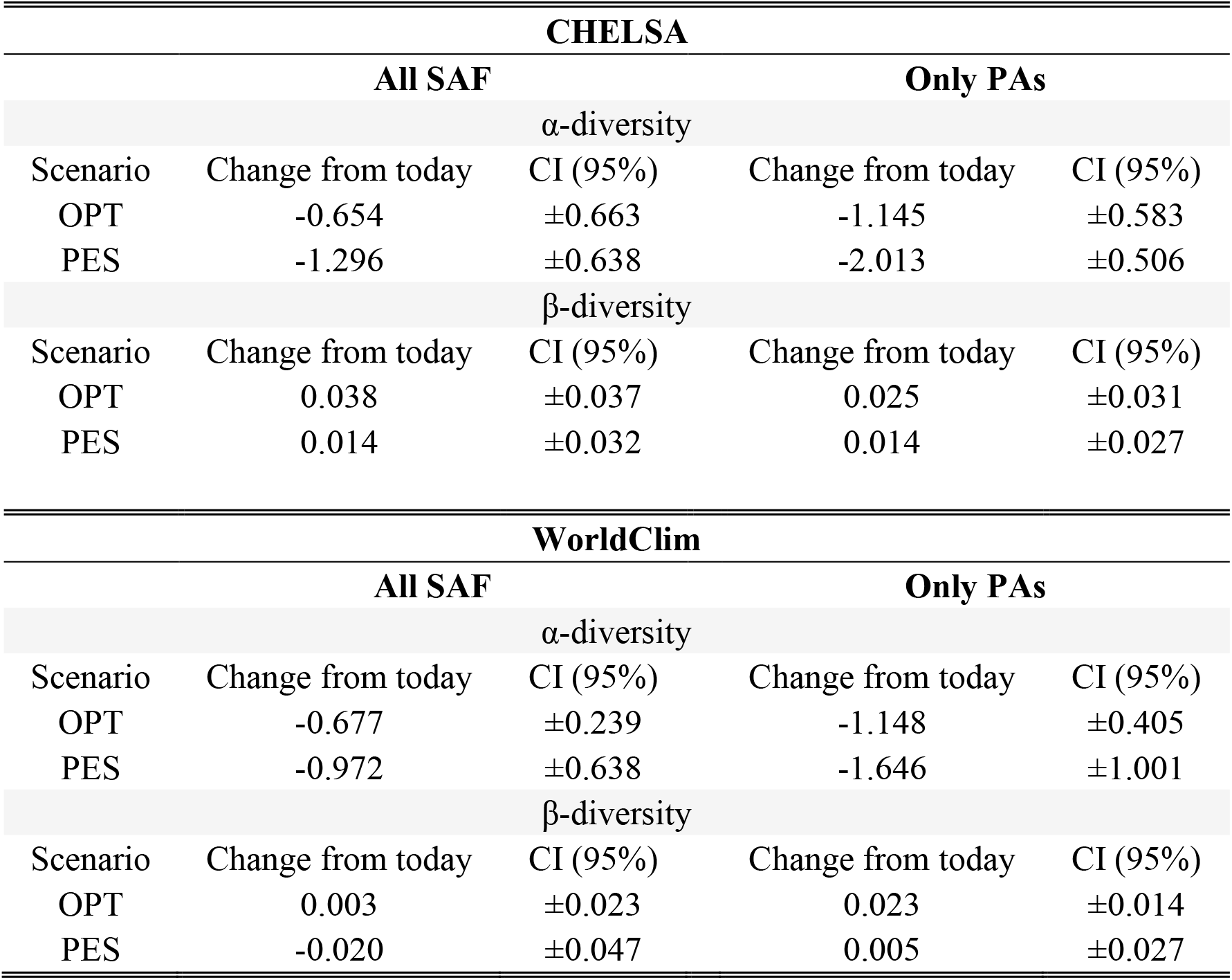
Regional changes in α - and β -diversity between each climate data set, scenario, and region. A significant change exists when the (absolute) predicted mean value is greater than the (absolute) confidence interval (CI). SAF – Subtropical Atlantic Forest; PA – Protected Areas only; OPT – optimistic future scenario; PES – pessimistic future scenario.

| CHELSA |  |  |  |  |
| --- | --- | --- | --- | --- |
| All SAF |  |  | Only PAs |  |
| α-diversity | | | | |
| Scenario | Change from today | CI (95%) | Change from today | CI (95%) |
| OPT | -0.654 | $\pm 0.663$ | -1.145 | $\pm 0.583$ |
| PES | -1.296 | $\pm 0.638$ | -2.013 | $\pm 0.506$ |
| β-diversity | | | | |
| Scenario | Change from today | CI (95%) | Change from today | CI (95%) |
| OPT | 0.038 | $\pm 0.037$ | 0.025 | $\pm 0.031$ |
| PES | 0.014 | $\pm 0.032$ | 0.014 | $\pm 0.027$ |

| WorldClim |  |  |  |  |
| --- | --- | --- | --- | --- |
| All SAF |  |  | Only PAs |  |
| α-diversity | | | | |
| Scenario | Change from today | CI (95%) | Change from today | CI (95%) |
| OPT | -0.677 | $\pm 0.239$ | -1.148 | $\pm 0.405$ |
| PES | -0.972 | $\pm 0.638$ | -1.646 | $\pm 1.001$ |
| β-diversity | | | | |
| Scenario | Change from today | CI (95%) | Change from today | CI (95%) |
| OPT | 0.003 | $\pm 0.023$ | 0.023 | $\pm 0.014$ |
| PES | -0.020 | $\pm 0.047$ | 0.005 | $\pm 0.027$ |

**Figure 2.**
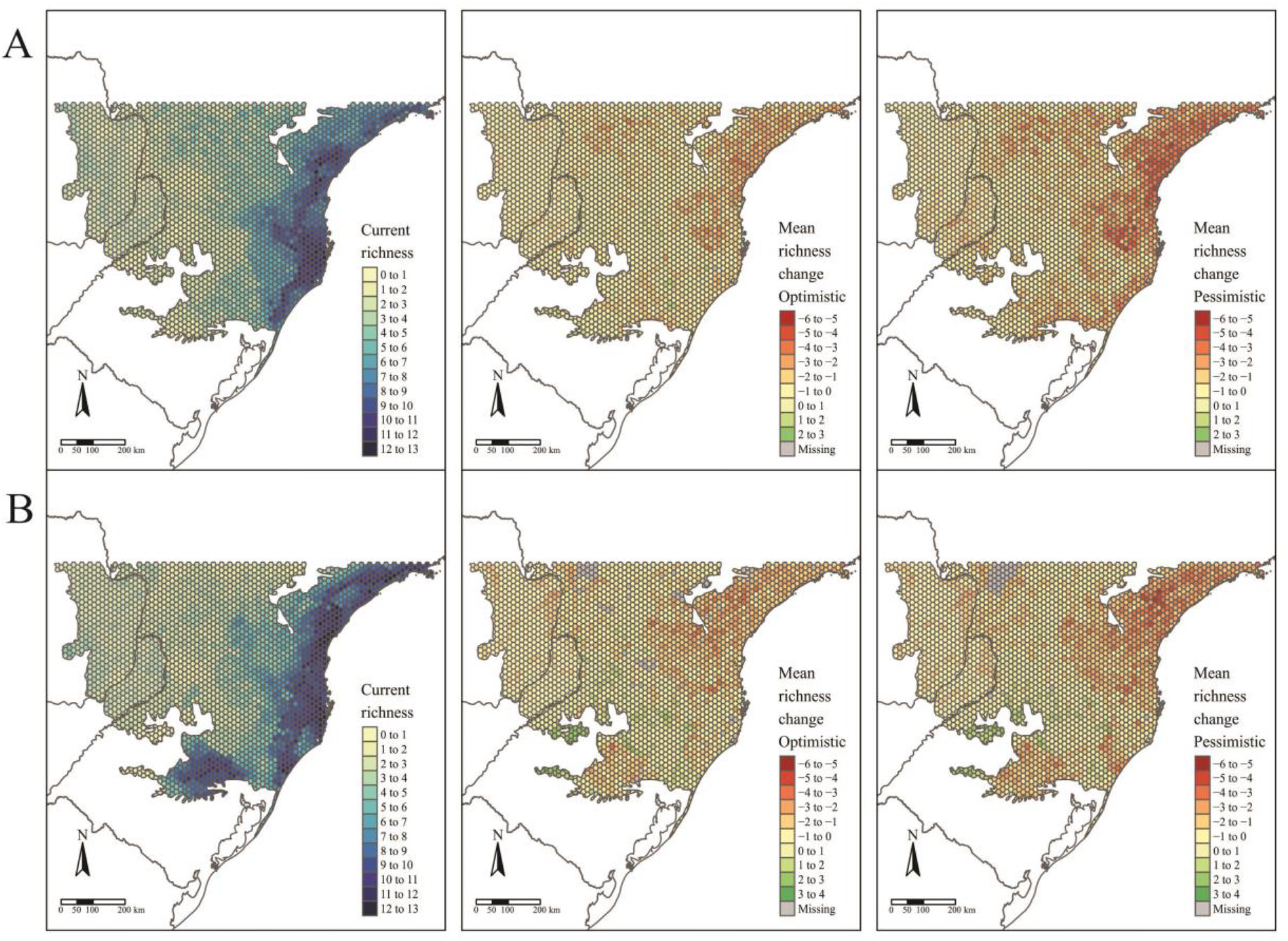
Patterns of tree ferns species richness in the subtropical Atlantic Forest in current time and the predicted change in the future scenarios using environmental data set from CHELSA (A) and WorldClim (B). The first-panel column shows current richness, the second-panel column shows the mean richness change in the optimistic scenario, and the third-panel column shows the mean richness change in the pessimistic scenario.

#### β-diversity

The regional BD_TOTAL_ in current time is 0.56 in CHELSA and 0.53 in WorldClim. In areas within PAs these values are lower: 0.50 and 0.46, respectively. In general, BD_TOTAL_ tends to not change in the future (Table 2). In CHELSA, BD_TOTAL_ tends to not change both future scenarios and inside PAs. The same patterns also take place in WorldClim — except in the pessimistic scenario inside PAs, where an increase in BD_TOTAL_ is predicted. LCBD presented similar patterns in both CHELSA and WorldClim predictions (Figure 3). In current time, higher LCBD values are detected in the north and west of the study area. These same areas are predicted to experience the larger changes in the future, losing their current uniqueness, i.e., less LCBD than in current time. Still, some areas in the Mixed Forest are predicted to increase its composition uniqueness.

**Figure 3.**
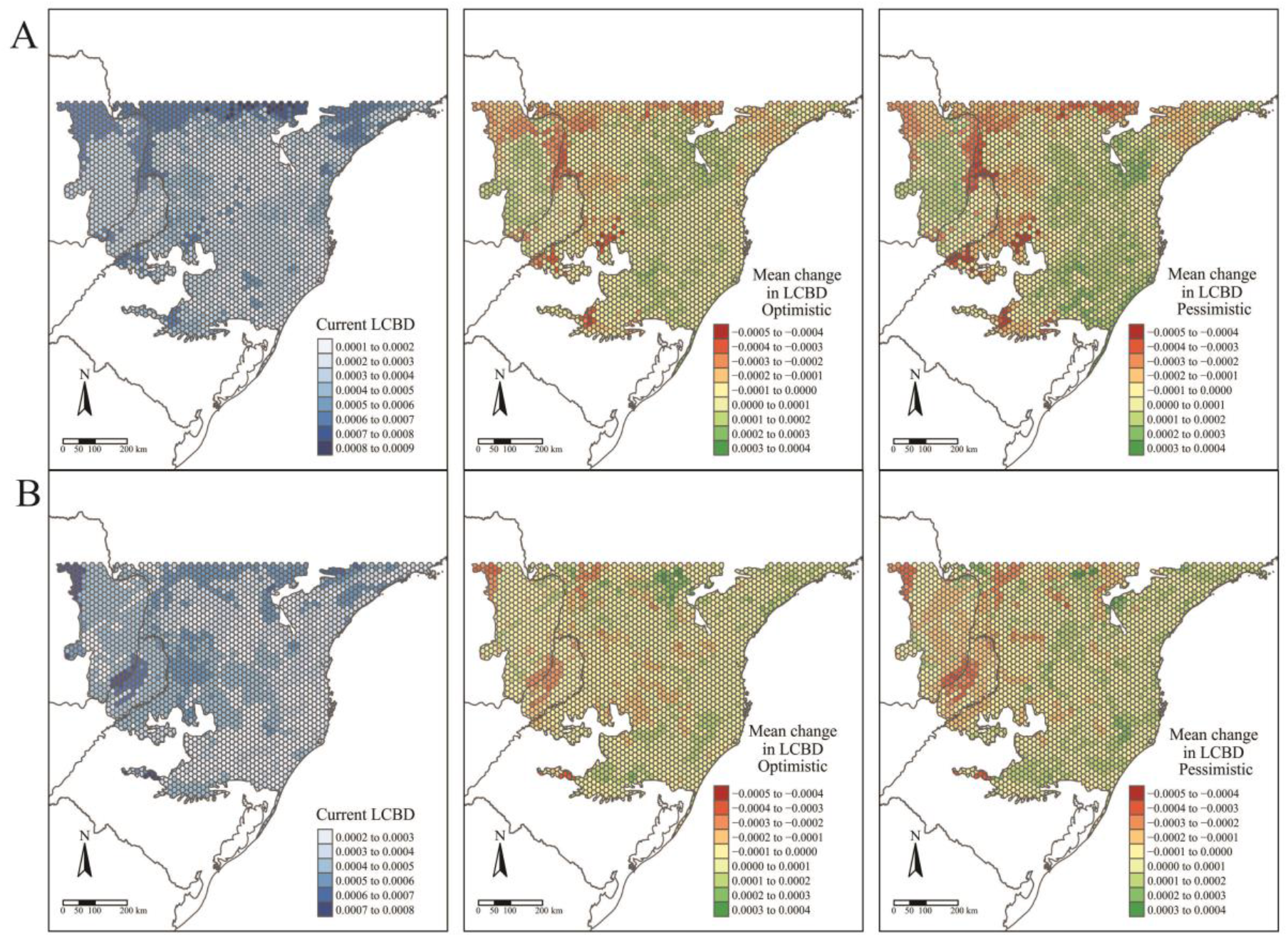
Patterns of tree ferns Local Contribution to β-diversity (LCBD) in the subtropical Atlantic Forest in current time and the predicted change in the future scenarios using environmental data set from CHELSA (A) and WorldClim (B). The first-panel column shows the current LCBD, the second-panel column shows the mean LCBD change in the optimistic scenario, and the third-panel column shows the mean LCBD change in the pessimistic scenario.

## DISCUSSION

Our first hypothesis (H_1_), which stated changes in species distribution due to climate changes was confirmed — despite our predictions being partially supported. Dicksoniaceae species indeed suffered a reduction in its distribution. However, contrary to our expectations, most of Cyatheaceae species also showed a decrease in their extents. Our second hypothesis (H_2_), that climate change will drive changes in species diversity (α- and β-diversity) on a regional scale — due to the impacts of species distribution change — was also partially supported. We found support for changes in species richness but not in BD_TOTAL_. Following the decreasing of species distribution in Cyatheaceae species, regional mean richness did not increase as we expected but decreased. On the regional scale, it seems that climate change will not affect β-diversity (BD_TOTAL_), but on a local scale (LCBD), the compositional uniqueness may change.

Our approach to measuring GCMs’ uncertainties was able to draw significant results for more than 70% of our combinations of climate data sets × regions × scenarios. This procedure preserved uncertainties and, seeing that we used the most interdependent GCMs (Sanderson *et al.* 2015), large intermodel variation was expected — expressed in CIs (Table 1, Figure S14-15). It is important to preserve (and account for) this source of variation since the choice of GCMs affect significantly the distribution of species (Diniz-Filho *et al.* 2009; Buisson *et al.* 2010).

### Changes in species distribution

A global analysis of Cyatheaceae distribution patterns showed a clear preference of the family for hotter and wetter locations with low seasonality — although some species show a capacity to occupy relatively cold regions, rarely where minimum temperatures drop below freezing (Bystriakova *et al.* 2011). Since an increase in temperature is expected, Cyatheaceae would be able to expand their range into parts of the Mixed Forest as the effects of higher temperatures could reduce the cold intensity (Wilson *et al.* 2019) and set up favorable conditions for some of Cyatheaceae species. Nonetheless, our results show that few Cyatheaceae species can expand their potential areas into the Semideciduous Forest, where there is a higher seasonality in precipitation than areas of Rainforest and eastern Mixed Forest (Oliveira-Filho *et al.* 2015). The two species that increased their distribution, *A. sternbergii* and *C. leucofolis*, are restricted within the northeast areas of the subtropics, probably with few populations having an austral distribution.

*Cyathea phalerata*, a common species in southern Brazil, is expected to lose distribution. This species thrives in warmer temperatures, *in vivo*, or *in vitro* (Marcon *et al.* 2017) — although temperatures above 32°C seem to inhibit spore germination. If this pattern recurs in all Cyatheaceae, a major increase in temperatures could restrict even more the species range in the future. *Alsophila setosa* — one of the most common tree ferns in forest communities within the subtropical Atlantic Forest (Lingner *et al.* 2015) — is also expected to lose distribution. This species can proliferate through vegetative growth and forest degradation can increase in density (Schmitt & Windisch 2005).

As we expected from Dicksoniaceae, both species tend to show an abrupt reduction in their distribution. For instance, in the pessimistic scenario, more than 65% of *D. sellowiana* distribution — an already endangered species — may be lost (more than 70% in *L*. *quadripinnata*). *Dicksonia sellowiana* is a species capable of withstanding colder environments (Gasper *et al.* 2011). For this reason, its future distribution tends to be restricted to higher and colder locations of the Mixed Forest (S01).

*Lophosoria quadripinnata* seems able to inhabit the colder regions of the Mixed Forest as well as part of the Rainforest. The species, just as *D. sellowiana*, also suffers a reduction in its distribution, being limited especially to the Mixed Forest. *Lophosoria quadripinnata* has been recorded in both temperate (Ricci 1996) and tropical (Bernabe *et al.* 1999) Rainforests. In this sense, precipitation — rather than temperature — appears to be the main determinant of this tree fern occurrence.

### Changes in diversity patterns

The second hypothesis was that distinctive responses of Dicksoniaceae and Cyatheaceae species should drive changes in α- and β-diversity patterns. We expected that species richness (α-diversity) would increase, following the increase in overlapping distributions of Cyatheaceae species which in turn would lead to a decrease of β-diversity — leading to diversity homogenization. Our expectations about α-diversity were not confirmed since a decrease in regional richness is predicted in the entire studied region and also on sites within PAs (Figure 2, Table 2). The current locations that uphold more richness are the ones that will be strongly affected by climate change. This summarizes most of the losses towards the north portion of the Atlantic Rainforest, the coastal region of our study area (Figure 1). These losses may affect the conservation of other biological groups that depend on tree ferns. For instance, some epiphytes have a substrate preference for them (Mehltreter 2008; Bartels & Chen 2012).

All observed changes may be associated with variations in precipitation since water availability seems to be an important species richness predictor for ferns, as pointed out by several authors (Aldasoro *et al.* 2004; Kessler *et al.* 2011; Gasper *et al.* 2015). Thus, the current low α-diversity of tree ferns observed in Mixed and Semideciduous forests, as well as the reduction of α-diversity in future scenarios, may be caused by rainfall regimes and higher climatic seasonality (Bystriakova *et al.* 2011; Cabré *et al.* 2016).

Changes in the β-diversity seem to be a little more complex. Although we did not find a trend in the changes in BD_TOTAL_, LCBD values point out changes in the relative uniqueness of species composition in some areas. In the west, a region with few species in the current climate and few losses in the future, LCBD is higher in the current climate and tends to decrease in the future. It seems that richness loss in the east region causes the composition of west sites to become more similar to other sites. Differently, the Mixed Forest region shows a tendency to become more unique, perhaps because richness becomes more similar to the other sites — while species composition differed due to its sites occurring within the center of a climatic gradient, i.e., in a transitional zone of species distribution.

### Implications for biodiversity conservation agenda

Our results provide relevant insights into the conservation of tree ferns by predicting which species will lose suitable habitats in the future (Table 1). Also, our results demonstrate a higher richness inside PAs than outside, indicating that the actual PAs are established in the richest areas of the subtropical Atlantic Forest (at least for tree ferns).

We note that almost all species of tree ferns will lose suitable areas inside the subtropical Atlantic Rainforest — not only Dicksoniaceae species as we thought. Species that occupy higher areas such as *A. capensis, L. quadripinnata*, and *D. sellowiana* seem to be exposed to more distribution constrains than species occurring in the Rainforest. Historically endangered species such as *D. sellowiana* had their potential distribution area greatly diminished. Also, climatic changes may alter their threatened status from endangered (Santiago *et al.* 2013) to critically endangered (considering IUCN 2012 criteria of population size reduction: A1cB1b).

Unfortunately, some PAs in higher altitudes, such as Parque Nacional de São Joaquim — where *D. sellowiana*, as well as other threatened species (such as *Araucaria angustifolia*; Wilson *et al.* 2019) may find suitable areas in the future — are threatened to be downsized (see the workgroup created to study the protected area boundaries; ICMBio – Instituto Chico Mendes de Conservação da Biodiversidade 2019).

More than half of the studied species are predicted to lose suitable areas, even inside PAs, where the losses are greater in some scenarios when compared to all subtropical regions (Table 1). So, to safeguard these species, not only new PAs will be needed, but the environmental legislation that protects the Atlantic Forest needs to be respected (conduct that does not exist in other Brazilian domains, such as the Amazon — Rajão *et al.* 2020). This includes protecting forest remnants inside particular properties, as they play an essential role in protecting species *in situ* (Chape *et al.* 2005; Metzger *et al.* 2019). Yet, new PAs may lose their long-term effectiveness in species conservation since they are also going to be impacted by climate change. To optimize cost-benefit analysis of implementing new PAs, lawmakers and specialists should always consider species conservation now and in the future (Araújo *et al.* 2011).

## Acknowledgments

The authors are thankful to Fundação de Amparo à Pesquisa e Inovação de Santa Catarina (FAPESC) for supporting IFFSC and for Coordenação de Aperfeiçoamento de Pessoal de Nível Superior - Brasil (CAPES) - Finance Code 001, for postgraduate research grants. We are also grateful for FUMDES (SC) and PIPE (SC, FURB) for funding research at the undergraduate level. The authors declare that they have no conflict of interest.

## Notes

https://avrodrigues.shinyapps.io/tferns/

### Competing Interest Statement

The authors have declared no competing interest.

### Summary of Updates

This version of the manuscript has been revised to update most of the methodological steps. Thus, this version amounts to new methods, results, and discussion.

## REFERENCES

Aldasoro, J.J., Cabezas, F. & Aedo, C. (2004) Diversity and distribution of ferns in sub-Saharan Africa, Madagascar and some islands of the South Atlantic. Journal of Biogeography 31, 1579–1604.

Alvares, C.A., Stape, J.L., Sentelhas, P.C., de Moraes Gonçalves, J.L. & Sparovek, G. (2013) Köppen’s climate classification map for Brazil. Meteorologische Zeitschrift 22, 711–728.

Araújo, M.B., Alagador, D., Cabeza, M., Nogués-Bravo, D. & Thuiller, W. (2011) Climate change threatens European conservation areas. Ecology Letters 14, 484–492.

Arens, N.C. & Baracaldo, P.S. (1998) Distribution of Tree Ferns (Cyatheaceae) across the Successional Mosaic in an Andean Cloud Forest, Nariño, Colombia. American Fern Journal 88, 60–71.

Arora, V.K., Scinocca, J.F., Boer, G.J., Christian, J.R., Denman, K.L., Flato, G.M., Kharin, V. V., Lee, W.G. & Merryfield, W.J. (2011) Carbon emission limits required to satisfy future representative concentration pathways of greenhouse gases. Geophysical Research Letters 38, 3–8.

Bartels, S.F. & Chen, H.Y.H. (2012) Mechanisms Regulating Epiphytic Plant Diversity. Critical Reviews in Plant Sciences 31, 391–400.

Bentsen, M., Bethke, I., Debernard, J.B., Iversen, T., Kirkevåg, A., Seland, Ø., Drange, H., Roelandt, C., Seierstad, I.A., Hoose, C. & Kristjánsson, J.E. (2013) The Norwegian Earth System Model, NorESM1-M – Part 1: Description and basic evaluation of the physical climate. Geoscientific Model Development 6, 687–720.

Bernabe, N., Williams-Linera, G. & Palacios-Rios, M. (1999) Tree Ferns in the Interior and at the Edge of a Mexican Cloud Forest Remnant: Spore Germination and Sporophyte Survival and Establishment. Biotropica 31, 83.

Boucher, O., Servonnat, J., Albright, A.L., Aumont, O., Balkanski, Y., Bastrikov, V., Bekki, S., Bonnet, R., Bony, S., Bopp, L., Braconnot, P., Brockmann, P., Cadule, P., Caubel, A., Cheruy, F., Codron, F., Cozic, A., Cugnet, D., D’Andrea, F., Davini, P., Lavergne, C., Denvil, S., Deshayes, J., Devilliers, M., Ducharne, A., Dufresne, J. -L., Dupont, E., Éthé, C., Fairhead, L., Falletti, L., Flavoni, S., Foujols, M. -A., Gardoll, S., Gastineau, G., Ghattas, J., Grandpeix, J. -Y., Guenet, B., Guez, L., Guilyardi, É., Guimberteau, M., Hauglustaine, D., Hourdin, F., Idelkadi, A., Joussaume, S., Kageyama, M., Khodri, M., Krinner, G., Lebas, N., Levavasseur, G., Lévy, C., Li, L., Lott, F., Lurton, T., Luyssaert, S., Madec, G., Madeleine, J. -B., Maignan, F., Marchand, M., Marti, O., Mellul, L., Meurdesoif, Y., Mignot, J., Musat, I., Ottlé, C., Peylin, P., Planton, Y., Polcher, J., Rio, C., Rochetin, N., Rousset, C., Sepulchre, P., Sima, A., Swingedouw, D., Thiéblemont, R., Traore, A.K., Vancoppenolle, M., Vial, J., Vialard, J., Viovy, N. & Vuichard, N. (2020) Presentation and evaluation of the IPSL-CM6A-LR climate model. Journal of Advances in Modeling Earth Systems, 1–52.

Brazilian Flora 2020 in construction (2020) Brazilian Flora 2020 in construction. Jardim Botânico do Rio de Janeiro.

Brock, J.M.R., Perry, G.L.W., Lee, W.G. & Burns, B.R. (2016) Tree fern ecology in New Zealand: A model for southern temperate rainforests. Forest Ecology and Management 375, 112–126.

Buisson, L., Thuiller, W., Casajus, N., Lek, S. & Grenouillet, G. (2010) Uncertainty in ensemble forecasting of species distribution. Global Change Biology 16, 1145–1157.

Burnham, K.P. & Anderson, D.R. (2002) 172 Ecological Modelling Model selection and multimodel inference: a practical information-theoretic approach. 2nd ed. Springer, New York.

Bystriakova, N., Schneider, H. & Coomes, D. (2011) Evolution of the climatic niche in scaly tree ferns (Cyatheaceae, Polypodiopsida). Botanical Journal of the Linnean Society 165, 1–19.

Cabré, M.F., Solman, S. & Núñez, M. (2016) Regional climate change scenarios over southern South America for future climate (2080-2099) using the MM5 Model. Mean, interannual variability and uncertainties. Atmósfera 29, 35–60.

Chape, S., Harrison, J., Spalding, M. & Lysenko, I. (2005) Measuring the extent and effectiveness of protected areas as an indicator for meeting global biodiversity targets. Philosophical transactions of the Royal Society of London. Series B, Biological sciences 360, 443–455.

Di Cola, V., Broennimann, O., Petitpierre, B., Breiner, F.T., D’amen, M., Randin, C., Engler, R., Pottier, J., Pio, D., Dubuis, A., Pellissier, L., Mateo, R.G., Hordijk, W., Salamin, N. & Guisan, A. (2017) ecospat: an R package to support spatial analyses and modeling of species niches and distributions. Ecography 40, 774–787.

Dee, D.P., Uppala, S.M., Simmons, A.J., Berrisford, P., Poli, P., Kobayashi, S., Andrae, U., Balmaseda, M.A., Balsamo, G., Bauer, P., Bechtold, P., Beljaars, A.C.M., van de Berg, L., Bidlot, J., Bormann, N., Delsol, C., Dragani, R., Fuentes, M., Geer, A.J., Haimberger, L., Healy, S.B., Hersbach, H., Hólm, E. V., Isaksen, L., Kållberg, P., Köhler, M., Matricardi, M., Mcnally, A.P., Monge-Sanz, B.M., Morcrette, J.J., Park, B.K., Peubey, C., de Rosnay, P., Tavolato, C., Thépaut, J.N. & Vitart, F. (2011) The ERA-Interim reanalysis: Configuration and performance of the data assimilation system. Quarterly Journal of the Royal Meteorological Society 137, 553–597.

Diniz-Filho, J.A.F., Mauricio Bini, L., Fernando Rangel, T., Loyola, R.D., Hof, C., Nogués-Bravo, D. & Araújo, M.B. (2009) Partitioning and mapping uncertainties in ensembles of forecasts of species turnover under climate change. Ecography 32, 897–906.

Dray, S., Bauman, D., Blanchet, G., Borcard, D., Clappe, S., Guénard, G., Jombart, T., Larocque, G., Legendre, P., Madi, N. & Wagner, H.H. (2019) adespatial: Multivariate multiscale spatial analysis. R package version 0.3–7.

Dunne, J.P., John, J.G., Adcroft, A.J., Griffies, S.M., Hallberg, R.W., Shevliakova, E., Stouffer, R.J., Cooke, W., Dunne, K.A., Harrison, M.J., Krasting, J.P., Malyshev, S.L., Milly, P.C.D., Phillipps, P.J., Sentman, L.T., Samuels, B.L., Spelman, M.J., Winton, M., Wittenberg, A.T. & Zadeh, N. (2012) GFDL’s ESM2 global coupled climate-carbon earth system models. Part I: Physical formulation and baseline simulation characteristics. Journal of Climate 25, 6646–6665.

Eisenlohr, P.V. & Oliveira-Filho, A.T. (2014) Tree species composition in areas of Atlantic Forest in southeastern Brazil is consistent with a new system for classifying the vegetation of South America. Acta Botanica Brasilica 28, 227–233.

Eleutério, A.A. & Pérez-Salicrup, D. (2006) Management of tree ferns (Cyathea spp.) for handicraft production in Cuetzalan, Mexico. Economic Botany 60, 182–186.

Fernandes, I. (1997) Taxonomia e fitogeografia de Cyatheaceae e Dicksoniaceae nas regiões Sul e Sudeste do Brasil. Universidade de São Paulo

Fick, S.E. & Hijmans, R.J. (2017) WorldClim 2: new 1-km spatial resolution climate surfaces for global land areas. International Journal of Climatology 37, 4302–4315.

Fielding, A.H. & Bell, J.F. (1997) A review of methods for the assessment of prediction errors in conservation presence/absence models. Environmental Conservation 24, 38–49.

Fourcade, Y., Besnard, A.G. & Secondi, J. (2018) Paintings predict the distribution of species, or the challenge of selecting environmental predictors and evaluation statistics. Global Ecology and Biogeography 27, 245–256.

Fourcade, Y., Engler, J.O., Rödder, D. & Secondi, J. (2014) Mapping species distributions with MAXENT using a geographically biased sample of presence data: A performance assessment of methods for correcting sampling bias. PLoS ONE 9, 1–13.

Galindo-Leal, C. & Câmara, I.G. (2005) Mata Atlântica : biodiversidade, ameaças e perspectivas. C. Galindo-Leal and I. de Gusmão Câmara (Eds). Fundação SOS Mata Atlântica — Belo Horizonte : Conservação Internacional.

Gasper, A.L. de, Eisenlohr, P.V. & Salino, A. (2015) Climate-related variables and geographic distance affect fern species composition across a vegetation gradient in a shrinking hotspot. Plant Ecology & Diversity 8, 25–35.

Gasper, A.L. de, Sevegnani, L., Vibrans, A.C., Uhlmann, A., Lingner, D.V., Verdi, M., Dreveck, S., Stival-Santos, A., Brogni, E., Schmitt, R. & Klemz, G. (2011) Inventário de Dicksonia sellowiana Hook. em Santa Catarina. Acta Botanica Brasilica 25, 776–784.

Giorgetta, M.A., Jungclaus, J., Reick, C.H., Legutke, S., Bader, J., Böttinger, M., Brovkin, V., Crueger, T., Esch, M., Fieg, K., Glushak, K., Gayler, V., Haak, H., Hollweg, H.-D., Ilyina, T., Kinne, S., Kornblueh, L., Matei, D., Mauritsen, T., Mikolajewicz, U., Mueller, W., Notz, D., Pithan, F., Raddatz, T., Rast, S., Redler, R., Roeckner, E., Schmidt, H., Schnur, R., Segschneider, J., Six, K.D., Stockhause, M., Timmreck, C., Wegner, J., Widmann, H., Wieners, K.-H., Claussen, M., Marotzke, J. & Stevens, B. (2013) Climate and carbon cycle changes from 1850 to 2100 in MPI-ESM simulations for the Coupled Model Intercomparison Project phase 5. Journal of Advances in Modeling Earth Systems 5, 572–597.

Hao, T., Elith, J., Lahoz-Monfort, J.J. & Guillera-Arroita, G. (2020) Testing whether ensemble modelling is advantageous for maximising predictive performance of species distribution models. Ecography 43, 549–558.

Hoshizaki, B.J. & Moran, R.C. (2001) Fern grower’s manual. Timber Press, Portland, Oregon.

Hurrell, J.W., Holland, M.M., Gent, P.R., Ghan, S., Kay, J.E., Kushner, P.J., Lamarque, J.F., Large, W.G., Lawrence, D., Lindsay, K., Lipscomb, W.H., Long, M.C., Mahowald, N., Marsh, D.R., Neale, R.B., Rasch, P., Vavrus, S., Vertenstein, M., Bader, D., Collins, W.D., Hack, J.J., Kiehl, J. & Marshall, S. (2013) The community earth system model: A framework for collaborative research. Bulletin of the American Meteorological Society 94, 1339–1360.

ICMBio – Instituto Chico Mendes de Conservação da Biodiversidade (2019) Portaria N°116, de 22 de março de 2019.

IPCC (2014) Climate Change 2014: Synthesis Report. Contribution of Working Groups I, II and III to the Fifth Assessment Report of the Intergovernmental Panel on Climate Change [Core Writing Team, R.K. Pachauri and L.A. Meyer (eds.)]. Geneva, Switzerland

Kaky, E., Nolan, V., Alatawi, A. & Gilbert, F. (2020) Ecological Informatics A comparison between Ensemble and MaxEnt species distribution modelling approaches for conservation : A case study with Egyptian medicinal plants. Ecological Informatics 60, 101150.

Karger, D.N., Conrad, O., Böhner, J., Kawohl, T., Kreft, H., Soria-Auza, R.W., Zimmermann, N.E., Linder, H.P. & Kessler, M. (2017) Climatologies at high resolution for the earth’s land surface areas. Scientific Data 4, 1–20.

Kessler, M., Kluge, J., Hemp, A. & Ohlemüller, R. (2011) A global comparative analysis of elevational species richness patterns of ferns. Global Ecology and Biogeography 20, 868–880.

Legendre, P. & De Cáceres, M. (2013) Beta diversity as the variance of community data: Dissimilarity coefficients and partitioning. Ecology Letters 16, 951–963.

Lehnert, M. & Kessler, M. (2018) Prodromus of a fern flora for Bolivia. XXI. Dicksoniaceae. Phytotaxa 344, 69.

Li, L., Lin, P., Yu, Y., Wang, B., Zhou, T., Liu, L., Liu, J., Bao, Q., Xu, S., Huang, W., Xia, K., Pu, Y., Dong, L., Shen, S., Liu, Y., Hu, N., Liu, M., Sun, W., Shi, X., Zheng, W., Wu, B., Song, M., Liu, H., Zhang, X., Wu, G., Xue, W., Huang, X., Yang, G., Song, Z. & Qiao, F. (2013) The flexible global ocean-atmosphere-land system model, Grid-point Version 2: FGOALS-g2. Advances in Atmospheric Sciences 30, 543–560.

Lima, A.A., Ribeiro, M.C., Grelle, C.E.V. & Pinto, M. (2019) Impacts of climate changes on spatio-temporal diversity patterns of Atlantic Forest primates. Perspectives in Ecology and Conservation 17, 50–56.

Lingner, D.V., Schorn, L.A., Sevegnani, L., de Gasper, A.L., Meyer, L. & Vibrans, A.C. (2015) Floresta Ombrófila Densa de Santa Catarina-Brasil: agrupamento e ordenação baseados em amostragem sistemática. Ciência Florestal 25, 933–946.

Marcon, C., Silveira, T., Schmitt, J.L. & Droste, A. (2017) Abiotic environmental conditions for germination and development of gametophytes of Cyathea phalerata Mart. (Cyatheaceae). Acta Botanica Brasilica 31, 0–0.

Martin, G.M., Bellouin, N., Collins, W.J., Culverwell, I.D., Halloran, P.R., Hardiman, S.C., Hinton, T.J., Jones, C.D., McDonald, R.E., McLaren, A.J., O’Connor, F.M., Roberts, M.J., Rodriguez, J.M., Woodward, S., Best, M.J., Brooks, M.E., Brown, A.R., Butchart, N., Dearden, C., Derbyshire, S.H., Dharssi, I., Doutriaux-Boucher, M., Edwards, J.M., Falloon, P.D., Gedney, N., Gray, L.J., Hewitt, H.T., Hobson, M., Huddleston, M.R., Hughes, J., Ineson, S., Ingram, W.J., James, P.M., Johns, T.C., Johnson, C.E., Jones, A., Jones, C.P., Joshi, M.M., Keen, A.B., Liddicoat, S., Lock, A.P., Maidens, A. V., Manners, J.C., Milton, S.F., Rae, J.G.L., Ridley, J.K., Sellar, A., Senior, C.A., Totterdell, I.J., Verhoef, A., Vidale, P.L. & Wiltshire, A. (2011) The HadGEM2 family of Met Office Unified Model climate configurations. Geoscientific Model Development 4, 723–757.

Mehltreter, K. (2008) Phenology and habitat specificity of tropical ferns. In: T. A. Ranker and C. H. Haufler (Eds), Biology and evolution of ferns and lycophytes. Cambridge University Press, Cambridge.

Metzger, J.P., Bustamante, M.M.C., Ferreira, J., Fernandes, G.W., Librán-Embid, F., Pillar, V.D., Prist, P.R., Rodrigues, R.R., Vieira, I.C.G. & Overbeck, G.E. (2019) Why Brazil needs its Legal Reserves. Perspectives in Ecology and Conservation 17, 91–103.

Muscarella, R., Galante, P.J., Soley-Guardia, M., Boria, R.A., Kass, J.M., Uriarte, M. & Anderson, R.P. (2014) ENMeval: An R package for conducting spatially independent evaluations and estimating optimal model complexity for Maxent ecological niche models. Methods in Ecology and Evolution 5, 1198–1205.

Naimi, B. (2015) usdm: Uncertainty analysis for species distribution models. R package version 1, 1–12.

Oliveira-Filho, A.T., Budke, J.C., Jarenkow, J.A., Eisenlohr, P.V. & Neves, D.R.M. (2015) Delving into the variations in tree species composition and richness across South American subtropical Atlantic and Pampean forests. Journal of Plant Ecology 8, 242–260.

Phillips, S.J., Anderson, R.P., Dudík, M., Schapire, R.E. & Blair, M.E. (2017) Opening the black box: an open-source release of Maxent. Ecography 40, 887–893.

Phillips, S.J., Anderson, R.P. & Schapire, R.E. (2006) Maximum entropy modeling of species geographic distributions. Ecological Modelling 190, 231–259.

Phillips, S.J., Dudík, M., Elith, J., Graham, C.H., Lehmann, A., Leathwick, J. & Ferrier, S. (2009) Sample selection bias and presence-only distribution models: implications for background and pseudo-absence data. Ecological Applications 19, 181–97.

Porfirio, L.L., Harris, R.M.B., Lefroy, E.C., Hugh, S., Gould, S.F., Lee, G., Bindoff, N.L. & Mackey, B. (2014) Improving the Use of Species Distribution Models in Conservation Planning and Management under Climate Change. PLOS ONE 9, e113749.

R Core Team (2020) R: a language and environment for statistical computing.

Radosavljevic, A. & Anderson, R.P. (2014) Making better MAXENT models of species distributions: Complexity, overfitting and evaluation. Journal of Biogeography 41, 629–643.

Rajão, R., Soares-Filho, B., Nunes, F., Börner, J., Machado, L., Assis, D., Oliveira, A., Pinto, L., Ribeiro, V., Rausch, L., Gibbs, H. & Figueira, D. (2020) The rotten apples of Brazil’s agribusiness. Science 369, 246–248.

Ribeiro, M.C., Metzger, J.P., Martensen, A.C., Ponzoni, F.J. & Hirota, M.M. (2009) The Brazilian Atlantic Forest: How much is left, and how is the remaining forest distributed? Implications for conservation. Biological Conservation 142, 1141–1153.

Ricci, M. (1996) Variation in distribution and abundance of the endemic flora of Juan Fernandez Islands, Chile Pteridophyta. Biodiversity and Conservation 5, 1521–1532.

Sanderson, B.M., Knutti, R. & Caldwell, P. (2015) A representative democracy to reduce interdependency in a multimodel ensemble. Journal of Climate 28, 5171–5194.

Santiago, A.C.P., Mynssen, C.M., Maurenza, D., Penedo, T.S.A. & Sfair, J.C. (2013) Dicksoniaceae. In: G. Martinelli and M. A. Moraes (Eds), Livro vermelho da flora do Brasil. Andrea Jakobsson: Instituto de Pesquisas Jardim Botânico do Rio de Janeiro, Rio de Janeiro, pp. 475–476.

Sarmiento, G., Pinillos, M. & Garay, I. (2005) Biomass variability in tropical american lowland rainforests. Ecotropicos 18, 1–20.

Scherrer, D., D’Amen, M., Fernandes, R.F., Mateo, R.G. & Guisan, A. (2018) How to best threshold and validate stacked species assemblages? Community optimisation might hold the answer. Methods in Ecology and Evolution 9, 2155–2166.

Schmitt, J.L. & Windisch, P.G. (2005) Aspectos ecológicos de Alsophila setosa Kaulf. (Cyatheaceae, Pteridophyta) no Rio Grande do Sul, Brasil. Acta Botanica Brasilica 19, 859–865.

Schwartz, C.E. & Gasper, A.L. de (2020) Environmental factors affect population structure of tree ferns in the Brazilian subtropical Atlantic Forest. Acta Botanica Brasilica 34, 204–213.

Swart, N.C., Cole, J.N.S., Kharin, V.V., Lazare, M., Scinocca, J.F., Gillett, N.P., Anstey, J., Arora, V., Christian, J.R., Hanna, S., Jiao, Y., Lee, W.G., Majaess, F., Saenko, O.A., Seiler, C., Seinen, C., Shao, A., Sigmond, M., Solheim, L., Von Salzen, K., Yang, D. & Winter, B. (2019) The Canadian Earth System Model version 5 (CanESM5.0.3). Geoscientific Model Development 12, 4823–4873.

Tatebe, H., Ogura, T., Nitta, T., Komuro, Y., Ogochi, K., Takemura, T., Sudo, K., Sekiguchi, M., Abe, M., Saito, F., Chikira, M., Watanabe, S., Mori, M., Hirota, N., Kawatani, Y., Mochizuki, T., Yoshimura, K., Takata, K., O’Ishi, R., Yamazaki, D., Suzuki, T., Kurogi, M., Kataoka, T., Watanabe, M. & Kimoto, M. (2019) Description and basic evaluation of simulated mean state, internal variability, and climate sensitivity in MIROC6. Geoscientific Model Development 12, 2727–2765.

Tryon, R.M. & Tryon, A.F. (1982) Ferns and allied plants: with special reference to tropical America. Springer New York, New York.

UNEP-WCMC (2019) Protected Area Profile for Latin America & Caribbean from the World Database of Protected Areas.

VanDerWal, J., Murphy, H.T., Kutt, A.S., Perkins, G.C., Bateman, B.L., Perry, J.J. & Reside, A.E. (2013) Focus on poleward shifts in species’ distribution underestimates the fingerprint of climate change. Nature Climate Change 3, 239–243.

Vibrans, A.C., Gasper, A.L. de, Moser, P., Oliveira, L.Z., Lingner, D.V. & Sevegnani, L. (2020) Insights from a large-scale inventory in the southern Brazilian Atlantic Forest. Scientia Agricola 77, e20180036.

Vibrans, A.C., Sevegnani, L., Lingner, D.V., Gasper, A.L. de & Sabbagh, S. (2010) Inventário florístico florestal de Santa Catarina (IFFSC): aspectos metodológicos e operacionais. Pesquisa Florestal Brasileira 30, 291–302.

Voldoire, A., Saint-Martin, D., Sénési, S., Decharme, B., Alias, A., Chevallier, M., Colin, J., Guérémy, J.F., Michou, M., Moine, M.P., Nabat, P., Roehrig, R., Salas y Mélia, D., Séférian, R., Valcke, S., Beau, I., Belamari, S., Berthet, S., Cassou, C., Cattiaux, J., Deshayes, J., Douville, H., Ethé, C., Franchistéguy, L., Geoffroy, O., Lévy, C., Madec, G., Meurdesoif, Y., Msadek, R., Ribes, A., Sanchez-Gomez, E., Terray, L. & Waldman, R. (2019) Evaluation of CMIP6 DECK Experiments With CNRM-CM6-1. Journal of Advances in Modeling Earth Systems 11, 2177–2213.

Voldoire, A., Sanchez-Gomez, E., Salas y Mélia, D., Decharme, B., Cassou, C., Sénési, S., Valcke, S., Beau, I., Alias, A., Chevallier, M., Déqué, M., Deshayes, J., Douville, H., Fernandez, E., Madec, G., Maisonnave, E., Moine, M.-P., Planton, S., Saint-Martin, D., Szopa, S., Tyteca, S., Alkama, R., Belamari, S., Braun, A., Coquart, L. & Chauvin, F. (2013) The CNRM-CM5.1 global climate model: description and basic evaluation. Climate Dynamics 40, 2091–2121.

Wagner, K., Mendieta-Leiva, G. & Zotz, G. (2015) Host specificity in vascular epiphytes: A review of methodology, empirical evidence and potential mechanisms. AoB PLANTS 7, 1–25.

Warren, D.L. & Seifert, S.N. (2011) Ecological niche modeling in Maxent: the importance of model complexity and the performance of model selection criteria. Ecological Applications 21, 335–342.

Warren, D.L., Wright, A.N., Seifert, S.N. & Shaffer, H.B. (2014) Incorporating model complexity and spatial sampling bias into ecological niche models of climate change risks faced by 90 California vertebrate species of concern. Diversity and Distributions 20, 334–343.

Watanabe, S., Hajima, T., Sudo, K., Nagashima, T., Takemura, T., Okajima, H., Nozawa, T., Kawase, H., Abe, M., Yokohata, T., Ise, T., Sato, H., Kato, E., Takata, K., Emori, S. & Kawamiya, M. (2011) MIROC-ESM 2010: model description and basic results of CMIP5-20c3m experiments. Geoscientific Model Development 4, 845–872.

Weigand, A. & Lehnert, M. (2016) The scaly tree ferns (Cyatheaceae-Polypodiopsida) of Brazil. Acta Botanica Brasilica 30, 336–350.

Wilson, O.J., Walters, R.J., Mayle, F.E., Lingner, D.V. & Vibrans, A.C. (2019) Cold spot microrefugia hold the key to survival for Brazil’s Critically Endangered Araucaria tree. Global Change Biology 25, 4339–4351.

Wu, T., Lu, Y., Fang, Y., Xin, X., Li, L., Li, W., Jie, W., Zhang, J., Liu, Y., Zhang, L., Zhang, F., Zhang, Y., Wu, F., Li, J., Chu, M., Wang, Z., Shi, X., Liu, X., Wei, M., Huang, A., Zhang, Y. & Liu, X. (2019) The Beijing Climate Center Climate System Model (BCC-CSM): The main progress from CMIP5 to CMIP6. Geoscientific Model Development 12, 1573–1600.

Yukimoto, S., Adachi, Y., Hosaka, M., Sakami, T., Yoshimura, H., Hirabara, M., Tanaka, T.Y., Shindo, E., Tsujino, H., Deushi, M., Mizuta, R., Yabu, S., Obata, A., Nakano, H., Koshiro, T., Ose, T. & Kitoh, A. (2012) A new global climate model of the Meteorological Research Institute: MRI-CGCM3: -Model description and basic performance-. Journal of the Meteorological Society of Japan 90, 23–64.

