## Supplementary material for "Expected impacts of climate change on tree ferns distribution and diversity patterns in subtropical Atlantic Forest": Table 1

**Table 1.** Mean changes (%) in the distribution of species regarding the climatic data set, future scenario, and region. Negative values represent predicted loss in species area and positive values represent predicted gains. Omitted values are non-significant changes. SAF – All subtropical Atlantic Forest; PA – Only Protected Areas; OPT – optimistic future scenario; PES – pessimistic future scenario.

| **SPECIES** | **CHELSA** | | | | **WorldClim** | | | |
| --- | --- | --- | --- | --- | --- | --- | --- | --- |
|  | **SAF** | | **PA** | | **SAF** | | **PA** | |
|  | **OPT** | **PES** | **OPT** | **PES** | **OPT** | **PES** | **OPT** | **PES** |
|  | Cyatheaceae | | | | | | | |
| *Alsophila setosa* | -38.31 | -54.51 | -41.13 | -60.83 | – | – | -20.49 | – |
| *Alsophila sternbergii* | 67.68 | 103.51 | 17.49 | 30.71 | 14.51 | – | 5.78 | – |
| *Cyathea atrovirens* | – | -39.74 | -27.90 | -39.64 | 24.55 | 39.03 | – | – |
| *Cyathea corcovadensis* | – | -46.57 | – | -48.38 | -27.63 | -34.21 | -20.96 | -36.68 |
| *Cyathea delgadii* | -40.63 | -43.80 | -57.47 | -75.40 | – | – | 10.24 | – |
| *Cyathea feeana* | – | – | – | – | -50.03 | -70.05 | -73.67 | -81.79 |
| *Cyathea hirsuta* | – | – | – | -20.39 | -12.89 | – | -2.11 | – |
| *Cyathea leucofolis* | 116.77 | 178.19 | 30.60 | 55.19 | 41.04 | 82.50 | 28.99 | – |
| *Cyathea phalerata* | – | -46.49 | -28.69 | -60.27 | -30.26 | -38.97 | -17.30 | – |
| *Gymnosphaera capensis* | -41.30 | -64.04 | -42.22 | -63.49 | -25.54 | -40.16 | -27.47 | -41.49 |
| *Sphaeropteris gardneri* | 137.33 | – | 138.11 | – | -41.14 | -62.21 | -68.45 | -78.66 |
|  | Dicksoniaceae | | | | | | | |
| *Dicksonia sellowiana* | -57.40 | -65.62 | -55.03 | -65.52 | – | -37.86 | – | -47.27 |
| *Lophosoria quadripinnata* | -64.47 | -74.08 | -60.98 | -71.56 | -48.94 | -57.91 | -63.51 | -67.48 |
