## Supplementary material for "Expected impacts of climate change on tree ferns distribution and diversity patterns in subtropical Atlantic Forest": Table 2

Table 2. Regional changes in α - and β -diversity between each climate data set, scenario, and region. A significant change exists when the (absolute) predicted mean value is greater than the (absolute) confidence interval (CI). SAF – Subtropical Atlantic Forest; PA – Protected Areas only; OPT – optimistic future scenario; PES – pessimistic future scenario.

| **CHELSA** | | | | | |
| --- | --- | --- | --- | --- | --- |
|  | **All SAF** | |  | **Only PAs** | |
| α-diversity | | | | | |
| Scenario | Change from today | CI (95%) |  | Change from today | CI (95%) |
| OPT | -0.654 | ±0.663 |  | -1.145 | ±0.583 |
| PES | -1.296 | ±0.638 |  | -2.013 | ±0.506 |
| β-diversity | | | | | |
| Scenario | Change from today | CI (95%) |  | Change from today | CI (95%) |
| OPT | 0.038 | ±0.037 |  | 0.025 | ±0.031 |
| PES | 0.014 | ±0.032 |  | 0.014 | ±0.027 |
| **WorldClim** | | | | | |
|  | **All SAF** | |  | **Only PAs** | |
| α-diversity | | | | | |
| Scenario | Change from today | CI (95%) |  | Change from today | CI (95%) |
| OPT | -0.677 | ±0.239 |  | -1.148 | ±0.405 |
| PES | -0.972 | ±0.638 |  | -1.646 | ±1.001 |
| β-diversity | | | | | |
| Scenario | Change from today | CI (95%) |  | Change from today | CI (95%) |
| OPT | 0.003 | ±0.023 |  | 0.023 | ±0.014 |
| PES | -0.020 | ±0.047 |  | 0.005 | ±0.027 |
